# Optimization of growth and electrosynthesis of PolyHydroxyAlcanoates by the thermophilic bacterium *Kyrpidia spormannii*

**DOI:** 10.1101/2021.10.25.465696

**Authors:** Guillaume Pillot, Soniya Sunny, Victoria Comes, Sven Kerzenmacher

## Abstract

The electrosynthesis of valuable compounds by biofilms on electrodes is being intensively studied since few years. However, so far, the actual biofilms growing on cathodes produce mainly small and relatively inexpensive compounds such as acetate or ethanol. Recently, a novel Knallgas bacterium, *Kyrpidia spormannii* EA-1 has been described to grow on cathodes under thermophilic and microaerophilic conditions, producing significant amounts of PolyHydroxyAlkanoates (PHAs). These PHA are promising sustainable bioplastic polymers with the potential to replace petroleum-derived plastics in a variety of applications. However, the effect of culture conditions and electrode properties on the growth of *K. spormannii* EA-1 biofilms and PHA production is still unclear.

In this study, we report on the optimization of growth and PHA production in liquid culture and on the cathode of a Microbial Electrosynthesis System. Optimization of the preculture allows to obtain high cell density of up to 8.5 Log10 cells·ml^-1^ in 48h, decreasing the time necessary by a factor of 2.5. With respect to cathodic biofilm formation, this study was focused on the optimization of three main operating parameters, which are the applied cathode potential, buffer pH, and the oxygen concentration in the feed gas. Maximum biofilm formation and PHA production was observed at an applied potential of −844mV *vs*. SCE, pH 6.5, O_2_ saturation of 2.5%. The PHA concentration in the biofilm reached a maximum of ≈26.8 μg·cm^-2^ after optimization, but at 2.9% the coulombic efficiency remains relatively low. We expect that further nutrient limitation will allow the accumulation of more PHA, based on a dense biofilm growth. In conclusion, these findings take microbial electrosynthesis of PHA a step forward towards practical implementation.

## INTRODUCTION

Microbial Electrosynthesis Systems (MES) are emerging technologies for the sustainable production of organic compounds and fuels. This technology is based on the ability of some microorganisms, called electrotrophs, to use electrons from the cathode of an electrochemical system as energy source, fixating CO_2_ into biomass and side products (Rabaey and Rozendal, 2010). Feeding the system with renewable electricity (solar panels, wind turbines etc.), and anthropogenic CO_2_ makes it a ground breaking concept to reduce the escalation of the current climatic situation (Lovley and Nevin, 2011). Since the first proof-of-concept, a decade ago, research has been carried out on the characterization and optimization of these processes (Prévoteau et al., 2020). Different aspects have been investigated - the biocatalyst, the conditions of culture, and the engineering of the system - in order to increase the productivity and the value of the molecules produced. Until recently, the technology was limited to the production of low added-value acetic acid by homoacetogenic biofilms, with low product concentration (≈ 12 g·L^-1^) (Vassilev et al., 2019) and low competitivity compared to fermentation processes. In the last years, the range of products has expanded, associating different metabolisms to elongate the carbon chain up to butyric or caproic acid, with product concentrations up to 3.2 g L^-1^ and 1.5 g L^-1^, respectively (Jourdin et al., 2018).

In order to increase the competitiveness of microbial electrosynthesis in comparison to classic fermentation, it is necessary to develop new biocatalysts able to produce high value-added compounds at high rate. Two strategies are been investigated, the engineering of novel metabolic pathways in already described electrotrophs (Kracke et al., 2018), or the isolation of new electrotrophs from the environment with interesting metabolic capabilities. The discovery of novel metabolisms requires to focus on extreme or unusual environments where microorganisms evolved in response of stresses by developing new metabolisms (Coker, 2016). Extreme conditions, such as high temperature, salinity, pressure or extreme pH are also profitable for MES operation (Jourdin and Burdyny, 2021). Indeed, the increase of optimal temperature is known to increase the metabolic rate of microorganisms, avoid contaminations and increase electrolyte conductivity. Higher salinity increases the conductivity of the electrolyte and general performances. The acidic or alkaline pH tolerances allow higher pH imbalance at the electrodes. Higher pressure increases CO_2_ solubility and availability. So far, only few extremophilic electrotrophs have been identified. Two acetogenic thermophiles, *Moorella thermoacetica* and *Moorella thermoautotrophica* were tested at temperatures up to 70°C (Faraghiparapari and Zengler, 2017). Pillot et al. (2020, 2021) have shown the enrichment of electrotrophic communities from deep-sea hydrothermal vents, dominated by *Archaeoglobales*, producing pyruvate, glycerol, and acetate at 80°C in seawater. These communities were dominated by *Archaeoglobales*, known to use the Wood–Ljungdahl pathway to fix CO_2_. Alqahtani et al. (2019) have shown the enrichment of halophilic homoacetogens in MES, dominated by *Marinobacter* sp., from Red Sea Brine Pool. Unfortunately, the metabolic ability of these electrotrophs doesn’t seems yet to increases the range of products or increases yields in a significant way. Recently, Reiner et al. (2020) have reported on a thermoacidophilic electrotrophic community enriched from geothermal hot springs on the Azores. From this community, they succeeded to isolate a novel microaerophilic Knallgas bacterium, *Kyrpidia spormannii* EA-1, able to produce PolyHydroxyAlcanoates (PHA) from CO_2_ on a cathode. Since this isolation, two additional strains were isolated from Pantelleria Island in Italy (Hogendoorn et al., 2020)

PHA are of great biotechnological interest as precursor for bioplastic production. They are bio-based and biodegradable polyesters, used as energy storage in intracellular granules or involved in maintenance of anoxic photosynthesis and sulfur cycle in microbial mats (Obruca et al., 2020). More than 150 different monomers can be combined leading to extremely different properties. Different species have been described to produce PHA, such as *Alcaligenes latus*, *Cupriavidus necator*, and *Pseudomonas putida*. PHA accumulation is usually produced by fermentation of feedstock and promoted when an essential nutrient for growth is present in limited amount in the cultivation medium, whereas an organic carbon source is in excess (Kourmentza et al., 2017). The actual production cost of PHA is still 3-4 times higher compared to conventional polymers such as polypropylene or polyethylene (Panuschka et al., 2019). Electrosynthesis of PHA could drastically reduce the production cost, by replacing costly organic carbon source by inexpensive CO_2_ and increasing the purity of the end product.

In this context, *Kyrpidia* strains are excellent candidates as biocatalyst for PHA electrosynthesis. However, prior to assess the competitivity of the process a significant effort of optimization is necessary. These microaerophilic organisms are highly sensitive to O_2_ concentration and experimental conditions can highly influence their growth rate and productivity. Previous cultivation methods of this strain required more than 7 days of liquid culture to obtain significant growth, slowing down its characterization and optimization in MES. In this study, we aimed to optimize the growth of *K. spormannii* EA-1 in liquid culture and on cathode for the production of PHA. For the optimization in liquid media, a protocol for the culture in serum bottle of another Knallgas bacteria, *Aquifex aeolicus* (Uzarraga et al., 2011), was adapted and optimized by experimental design on different factors: the quantity of gaseous substrate represented by the ratio gas/liquid, the redox state of the medium (anaerobic/aerobic preparation), the mixing during incubation and the electron donor nature. For the electrotrophic growth on cathode, the electrode potential, the oxygen concentration and the pH of the media were tested independently and the associated current consumption, biofilm produced, and accumulation of PHA were quantified.

## MATERIAL AND METHODS

### Bacterial strain and culture media

*K. spormannii* EA-1 cultures were obtained from cryostock from the Applied Biology group of Johannes Gescher at the Karlsruhe Institute of Technology (Germany) and sub-cultured at 4% in 100ml serum bottles closed with a rubber stopper and filled with ES-medium before inoculation of the experimental design media or Microbial Electrochemical Systems (MES). The ES medium was prepared anaerobically (medium with low redox potential, coded N_2_) or not (medium with high redox potential, coded O_2_), with the following content (all procured from Carl Roth, Germany) per litre: 0.53 g NH_4_Cl, 0.15 g of NaCl, 0.04 g of KH_2_PO_4_, 0.2 g of yeast extract, 1 ml of 0.1 M CaCl_2_, 0.12 ml of 1 M MgSO_4_, and 1 ml of Wolfe’s Mineral Elixir (Wolin et al., 1963) and was adjusted at pH 5.5. In the anaerobic preparation, the media was supplemented with 0.5 g/L of Cysteine-HCl and 1mg/L of Resazurin, boiled for 15min and cooled down under N_2_ degassing. The volume in the serum bottles was adjusted at 50ml media + 60ml gas head space or 25ml media + 85ml gas head space and autoclaved. After inoculation, the headspace of the serum bottle was replaced by air at atmospheric pressure, and as electron donor either 20mM of Acetate a mixture of H_2_:CO_2_ (80:20) at an overpressure of 1.5 bar were added for the liquid cultures, depending on the condition tested. The media were incubated at 60°C and either shaken at 150 rpm in an incubator (Incubator 3032, GFL, Germany) or kept static (Incudrive H, Schuett Biotec, Germany).

### Experimental design for the optimization of liquid culture growth

The software Design Expert v13.0 was used to perform a two-level factorial design with four factors: shaking of the media, substrate used, redox state of the medium, and ratio gas/liquid. The conditions for the 16 runs are presented Table 1. Three responses were measured each day for 3 days to evaluate the growth of *K. spormannii*: optical density (OD) at 600nm, qPCR quantification, and PHA quantification (see below for details).

**Table 1:** Experimental Design for the optimization of the growth of *Kyrpidia spormannii* in liquid media on the four factors with A: Shaking, B: Substrate, C: Media preparation, D: Ratio liquid/gas in the bottle.

|  | A | B | C | D |
| --- | --- | --- | --- | --- |
|  | Shaking | Substrate | Media preparation | Ratio liquid/gas |
| Run 1 | Yes | H<sub>2</sub>:CO<sub>2</sub> | N<sub>2</sub> | 50/60 |
| Run 2 | Yes | H<sub>2</sub>:CO<sub>2</sub> | O<sub>2</sub> | 25/85 |
| Run 3 | Yes | Acetate | O<sub>2</sub> | 50/60 |
| Run 4 | No | H<sub>2</sub>:CO<sub>2</sub> | O<sub>2</sub> | 50/60 |
| Run 5 | No | Acetate | O<sub>2</sub> | 50/60 |
| Run 6 | No | Acetate | N<sub>2</sub> | 50/60 |
| Run 7 | No | Acetate | N<sub>2</sub> | 25/85 |
| Run 8 | No | H<sub>2</sub>:CO<sub>2</sub> | N<sub>2</sub> | 25/85 |
| Run 9 | No | Acetate | O<sub>2</sub> | 25/85 |
| Run 10 | Yes | H<sub>2</sub>:CO<sub>2</sub> | O<sub>2</sub> | 50/60 |
| Run 11 | No | H<sub>2</sub>:CO<sub>2</sub> | N<sub>2</sub> | 50/60 |
| Run 12 | Yes | Acetate | N<sub>2</sub> | 50/60 |
| Run 13 | No | H<sub>2</sub>:CO<sub>2</sub> | O<sub>2</sub> | 25/85 |
| Run 14 | Yes | Acetate | O<sub>2</sub> | 25/85 |
| Run 15 | Yes | Acetate | N<sub>2</sub> | 25/85 |
| Run 16 | Yes | H<sub>2</sub>:CO<sub>2</sub> | N<sub>2</sub> | 25/85 |

Each run was performed as triplicate and the average of each response was used during the ANOVA test. The selection of the factors was performed on the full factor interactions with the auto-selection using the AICc criteria and respecting the hierarchy. The OD_600nm_ measurement was performed on a spectrophotometer and the OD_600nm_ at 24h, 48h and 72h was normalized to the OD_600nm_ after inoculation.

### Microbial Electrochemical System for electrotrophic growth experiments

Optimization of biofilm growth was performed in a 6-electrode battery glass reactor, previously described (Erben et al., 2021), for the optimization of cathode potential, and in H-cells to allow the separation of conditions for optimization with respect to O_2_ concentration and pH. The cathode was a 2.25 cm^2^ exposed surface of graphite plate (Müller & Rössner GmbH & Co KG, Germany), the anode was a Ir-Ta mesh (Umicore, Belgium; ~15×15 mm), and the reference electrode was a Saturated Calomel Electrode (SCE, offset of −215mV vs. SHE at 60°C, Sensortechnik Meinsberg, Germany). The cathodes were rinsed with DI water and cleaned in an ultrasonic bath for 5 mins prior to be connected to a potentiostat (IPS Elektroniklabor, PGU-MOD-500mA, Münster, Germany) by titanium wires. The media was filled in the systems, 0.1M of PBS buffer with required pH was added, then the systems were closed and autoclaved. The systems were agitated with a magnetic stirrer at 150 rpm. The gas mixture (N_2_:CO_2_ at 77.5:20) was purged continuously in the system using flow meters (Analyt-MTC, Germany) and the O_2_ concentration was adjusted and monitored by an oxy-meter (Oxy-4 Mini, PreSens, Germany). The MES were placed in an incubator (Schuett Biotec.de, Incudrive H, Germany) at a constant temperature of 60°C. When the conditions were stabilized after 4 h, the system was inoculated at 2%(v/v) with a liquid culture obtained after 48h with the optimal conditions identified in the previous part.

### Fluorescence microscopy

Fluorescence microscopy analysis was used for visual confirmation and quantification of biofilm formation on the cathode after the electrochemical experiments. Upon completion of the experiment, bacterial cells were fixed to the electrode using 4% glutaraldehyde in PBS 0.1M for 30 mins and later washed in DI water. The fixed electrodes were stained with 2 μg·ml^-1^ DAPI (4’,6-diamidino-2-phenylindole) and Nile Red (Carl Roth, Germany) and incubated in the dark for 30 mins. The stained biofilm on the electrode material were visualised using a Zeiss Microscope Axioscope 5/7 (Solid-State Light source Colibri 3 (Type RGB-UV), Microscopy Camera Axiocam 702 mono) (Zeiss, Germany) at 250x magnification (Objective ApoChrom 25x) under oil immersion and subsequently the z-stacks were automatically captured with the motorized stage on the Zen software (Zeiss, Germany, version 3.0). The fluorescence microscopy image data were further processed to obtain the Z projection of the image stacks and the cell counting was done using Cellc12 software (Selinummi et al., 2005)

### Biofilm quantification by means of qPCR

After the experiment, the cathodes containing the biofilm were taken from the bioelectrochemical reactor and sonicated for 10 min in 10 ml DI water in order to detach the biofilm from the electrode surface. Furthermore, the 16S rRNA gene was partially amplified by the qPCR method in an Eco 48 Real Time PCR System (PCRmax, United Kingdom), using the qPCRBio SyGreen 2x-Mix (Nippon Genetics Europe, Germany), and the primers Alyc630F (5’-GAGAGGCAAGGGGAATTCC-3’) and 806R (5’-GGACTACHVGGGTWTCTAAT-3’). A standard curve was prepared through cloning method using pGEM(R)-T Easy Vector System II (Promega) and JM109 Competent Cells (Promega). The plasmid was extracted with PureYield™ Plasmid Miniprep System (Promega) and quantified on a Quantus™ Fluorometer using the QuantiFluor(R) dsDNA System (Promega). The quantification of copies of 16S rDNA was divided by the number of copies naturally present per cell (5 copies·cell^-1^ according to rrnDB database), to obtain the number of cells·ml^-1^.

### PHA quantification

Sonicated biofilm samples (see above) were prepared by alkaline hydrolysis according to Watanabe et al., (2012) with 1 ml of sample in 500 μl of 3N NaOH, heated at 100°C for 3 h, then neutralized with 500 μl of 3M HCl. A standard solution of poly(3-hydroxy-butyrate) (average Mn ~500,000, Sigma Aldrich) was prepared using the same method. A HPLC system (Alliance, Waters) equipped with a UV/Vis detector (2489 Detector, Waters) monitored at 210 nm was used for the analysis of crotonic acid produced by the hydrolysis step. The column was a Waters Atlantis C18, (Waters, United Kingdom, 250 mm × 4.5 mm, particle size 5 μm). The column temperature was set to 30 °C. The mobile phase was 0.014 N H_2_SO_4_ at a flow rate of 0.7 mL min^-1^. Before injection, samples were filtered through 0.45 μm pore size membrane filter (Minisart High Flow, Sartorius, Germany). A volume of 10 μL was injected into the instrument for analysis.

### Coulombic Efficiency

The Coulombic efficiency (CE) of the PHA production was calculated using the formula:

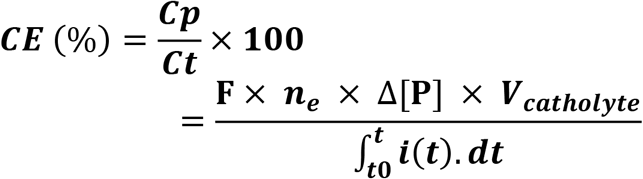

*C_T_*: total coulombs consumed
*C_P_*: coulombs found in the product
n_e_: Number of mol of electrons per mol of product
*F*: Faradays constant (96.485 C/mol)
Δ[P]: Variation of product concentration from t0 to tot
*V_catholyte_*: Volume of reaction

The total amount of current consumed by the system was calculated by integrating the area under current (A) vs. time (s). The quantity of electrons contained in the final product was calculated using 66 e^-^ equivalent per mole of PHB (Islam Mozumder et al., 2015), obtained from the stoichiometry of PHA production in autotrophic condition, using 33 moles of H_2_ for the production of 1 monomer of PHA.

## RESULTS AND DISCUSSION

### Optimization of growth of Kypridia spormannii in liquid media

The effect of four factors on the growth of *K. spormannii* EA-1 in liquid media was tested: The shaking of the bottle, the substrate used, the media preparation (aerobic, anaerobic, addition of reducing agent) and the gas/liquid volume ratio. These four factors directly or indirectly affect the oxygen concentration, which can be limiting or toxic (O_2_ quantity and transfer), the growth kinetics (H_2_ or Acetate), and the redox state of the medium (resazurin, H_2_, O_2_). The growth was measured by three techniques: OD_600nm_ determination, qPCR quantification, and PHA production. The respective graphs are presented in Supplementary data S1. The ANOVA analysis of the experimental design on the 3 responses are presented in Table 2.

**Table 2:**
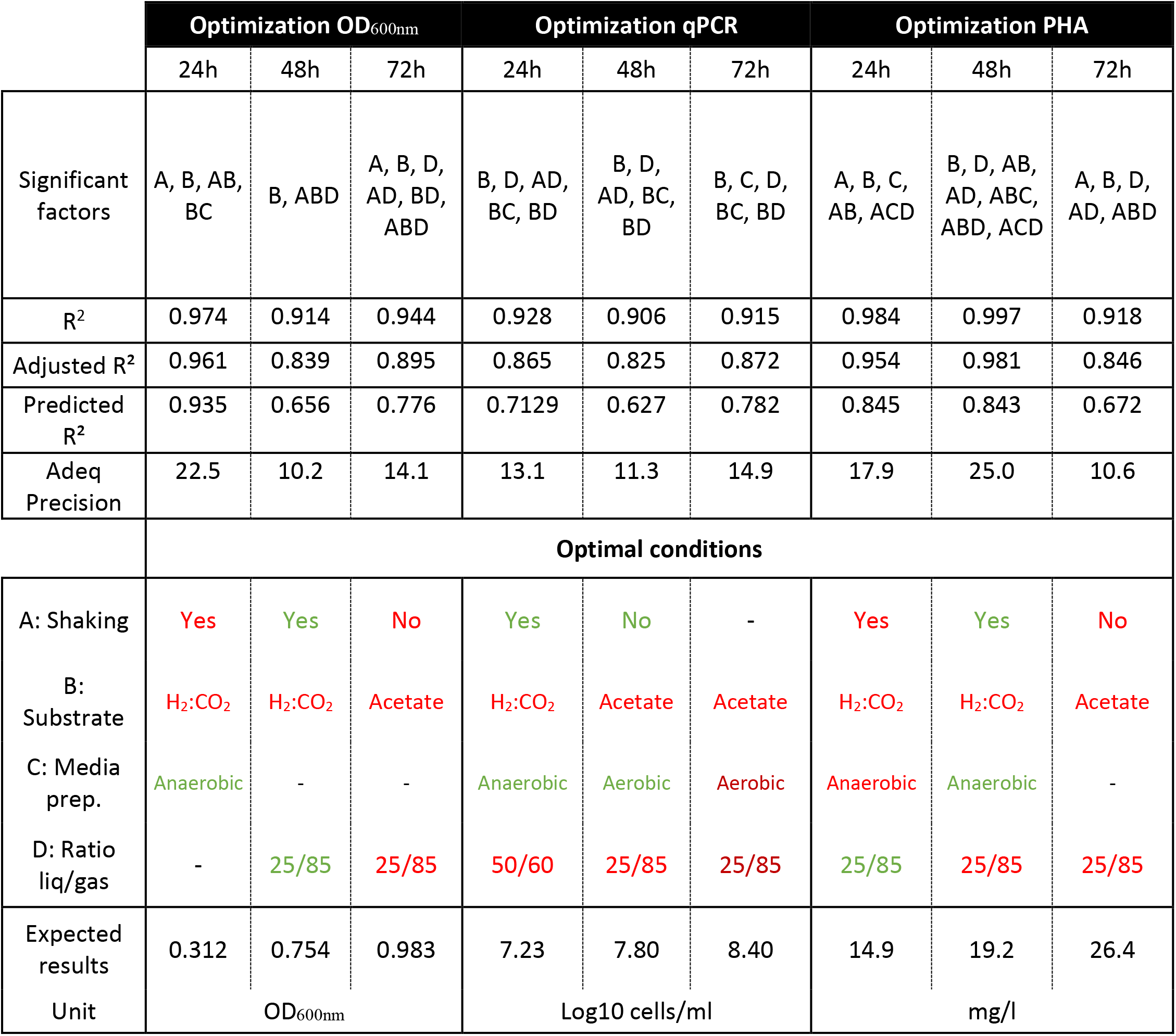
Results of the ANOVA analysis obtained according to the experimental design with factors A: Shaking, B: Substrate, C: Media preparation, D: Ratio liquid/gas in the bottle. Three responses - the OD_600nm_, the qPCR quantification and the PHA measurement - were considered at 3 times post inoculation on the average of triplicate experiments. The factors in red are main factors shown significant by the ANOVA test in the determination of the expected response. The factors in green are factors with a significant effect on the response when in interaction with other factors. The optimal conditions shown were defined by the ANOVA to maximize each response

As shown in Fig. S1, in most experimental runs the OD_600nm_ increased, a maximum of 0.993 ± 0.09 was observed in case of run 7. Only the Run 14 and 6 didn’t show any growth during the 4 days of experiment. The runs 7 and 15 showed a delay in the growth with a plateau only at 3 days, while all others plateaued after 2 days. The coefficient of determination R^2^ obtained by the ANOVA model for 24 h, 48 h, and 72 h were all relatively high, with a Predicted R^2^ in reasonable agreement with the Adjusted R^2^ (difference is less than 0.2), and the Adequate Precision is greater than 4, indicating that the ANOVA model is significantly representative. It allows to identify that the shaking of the media and the use of H_2_:CO_2_ instead of acetate had a significant positive effect on the OD_600nm_ after 24 h, to obtain a maximum OD_600nm_ of 0.312. The media preparation method had only an effect in combination with the substrate used. After 48 h, the use of H_2_:CO_2_, with anaerobic media preparation and a ratio 25 ml media/85 ml gas had a positive effect on the OD_600nm_ compared to their respective alternatives, yielding a maximum of 0.754. The shaking presented a significant effect only in combination with substrate and ratio factors.

Finally, when assessed after 72 h, only the media preparation didn’t have a significant effect on growth. A maximum OD_600nm_ of 0.983 was achieved with static culturing with acetate and a ratio of 25 ml liquid and 80 ml gas. These results indicate that a faster growth is obtained during the first day with H_2_ as substrate, to then reach a limitation with O_2_ concentration after 48 h and subsequently reach a plateau with most of conditions after 72 h. The OD_600nm_ measurement is a quick technique to assess the growth of most microorganisms but can be falsified by the production of intracellular granules or EPS, increasing artificially the absorbance with a constant number of cells. To overcome this potential issue, a second quantitative method was performed, based on the quantification of the 16S rDNA by qPCR.

The qPCR measurement shows a cell concentration (corrected with the number of copies of 16S rDNA per cells) of 5.32 ± 0.22 Log_10_cells·ml^-1^ after inoculation, increasing up to a maximum of 8.43 ± 0.56 Log_10_cells·ml^-1^ in the run 9 after 72 h. Only Run 6 didn’t show any growth, with slight growth on Run 14, that was not visible on the OD_600nm_, potentially due to a higher detection threshold with OD_600nm_ measurement or heterogeneity in the samples. Most of the runs with acetate (3, 5, 6, 12, 14 and 15) showed lower growth than the runs with H_2_:CO_2_. The fit statistics of the ANOVA indicated that all 3 models were significant (Table 2). The model shows higher cell density with H_2_:CO_2_ and a gas/liquid ratio of 50/60 after the first day, and with acetate and a ratio of 25/85 at 48 h and 72 h. The maximum cell concentrations in the identified optimal conditions are 7.23, 7.80 and 8.40 Log_10_cells·ml^-1^ at 24 h, 48 h and 72 h respectively. The difference between the qPCR and OD_600nm_ results, with poor correlations (maximum of R^2^=0.766 at 48 h) presented Fig S2-A, could be explained by the production of PHA over the growth, only detected with OD_600nm_ measurements.

The PHA quantification shows an increase from 3.0 ± 0.3mg·L^-1^ to up to 29.3 ± 1.2 mg·L^-1^ on run 7 after 72 h. The coefficients of determination at 24 h and 48 h are close to 1 but decrease to 0.83 after 72 h. The statistics of the ANOVA models show a good fit to our data. During the first days, the shaking, the use of H_2_:CO_2_, the anaerobic media preparation in combination with a volume ratio of 25/85 are significant factors on the PHA production, allowing to reach maximums of 14.9 mg·L^-1^ at 24 h and 18.8 mg·L^-1^ at 48 h. After 72 h, the use of acetate in a static culture became the best conditions to reach a maximum PHA production of 26.3 mg·L^-1^. As previously observed on OD_600nm_, the use of H_2_:CO_2_ as substrate and a good mixing allow a faster growth and PHA production, but additionally, the presence of a reduced media seems to induce the production of PHA. The higher PHA production in the absence of shaking after 72 h could also be explained by the lower O_2_ dissolution into the liquid. Indeed, in *Cupriavidus necator*, it was reported that O_2_ limitation enhance the PHA production, as energy storage, until the cells retrieve more favourable conditions (Kourmentza et al., 2017).

After 72 h of culture, most of the conditions reached a plateau or a decline, with high growth (Figure S1A), which is a net improvement from the previous culturing method requiring more than 7 days. Interestingly, our result seems to indicate a faster growth on H_2_ than on acetate, while the Gibbs free energy of the reaction of acetate oxidation release more energy (ΔG^0^ Acetate/O_2_ = −882 kJ mol^-1^ at 60°C) than the hydrogen oxidation (ΔG^0^ H_2_/O_2_ = −261 kJ mol^-1^ at 60°C) (Amend and Shock, 2001). However, it is known that acetate needs an activation step by the Acetyl-coenzyme A synthetase, that catalyzes the ATP- and CoA-dependent activation of acetate generating acetyl-CoA, AMP and pyrophosphate (acetate + ATP + CoA → acetyl-CoA + AMP + PP_i_) prior to enter the metabolism (Reiner et al., 2018b). In the hydrogenotrophic pathway, the H_2_ is directly converted into H^+^, used further by the ATP synthase to produce ATP (Brigham, 2019). This initial ATP consumption for acetate can explain the lag-time before growth in this condition. Similar results were observed in strains FAVT5 and COOX1, with doubling times of 3.6 h on H_2_ and 6 h on Acetate (Hogendoorn et al., 2020).

To better understand the effect of the media redox state and the ratio liquid/gas, the concentration of oxygen available in the serum bottles was calculated at 25°C, which is the temperature of media preparation and inoculation. The dissolved oxygen during aerobic media preparation plays a minor role in the total oxygen amount, as only 6.4 and 12.8 μmol of oxygen are present in 25 ml and 50 ml of media, respectively. However, the air flushed in the headspace of the bottle after autoclaving brings 0.799 mmol and 0.564 mmol of oxygen when the bottle is filled with 25 ml and 50 ml of media, respectively. On the other hand, the reducing agent added to the anaerobic media preparation (Cystein-HCl), and in a minor part the redox indicator resazurin, will react with O_2_ and remove up to 0.079 and 0.160 mmol in the 25 ml and 50 ml media, respectively.

The total H_2_ reaches 4.46 and 3.15 mmol with 25 ml and 50 ml of media, respectively. Considering the stoichiometry of already reported *Kyrpidia* strains of 1 mole of H_2_ for 0.36 mole of O_2_, the availability of O_2_ is limiting in our condition (Hogendoorn et al., 2020). As the oxygen sensitivity of *K. spormannii* EA-1 has not been evaluated yet, this difference of concentration can affect the growth significantly. *Aquifex aeolicus*, another microaerophilic (hyper)thermophilic bacteria, can grow with O_2_ concentration as low as 7.5 ppm (Deckert et al., 1998). Moreover, the volume ratio and the shaking influence the oxygen and hydrogen transfer to the liquid media during their consumption.

As previously mentioned, the difference of growth quantification by OD_600nm_ measurement and by qPCR can be explained by the absorbance of PHA at 600 nm. Figure S2-B represents the correlation between the OD_600nm_ and the PHA measurement for the 3 different sampling times. At t0, a poor correlation is observed, with R^2^ at 0.02, but increase quickly above 0.81 after 24 h, with a ratio converging to 28.4 ± 6.78 mg·L^-1^ of PHA per OD unit. Poorer correlations were observed between qPCR measurements and PHA quantification (Fig S2-C), with R^2^ at 0.22, 0.47, 0.79 and 0.70 on samples after inoculation, 24 h, 48 h, and 72 h respectively. The average ratio PHA/qPCR were 16.0, 10.1, 6.9 and 4.1 μg·cell^-1^, at 0 h, 24 h, 48 h and 72 h respectively, indicating a divergence of energy into cell multiplication rather than to PHA production during the course of the culture.

Concerning the PHA production, Kourmentza et al. (2017) report PHA concentrations between 0.08 to 2.7 g·L^-1^ (based on reactor volume) produced by different strains using organic carbon sources. Comparatively, our production of PHA is relatively low (maximum of 29.3 ± 1.2 mg·L^-1^), and could probably be optimized by nutrient limitation, as previously described. Up to 90% of the dry cell mass can be composed of PHA (Verlinden et al., 2007). In our case, assuming a mass of 10^-12^ g·cell^-1^, we could theoretically reach 0.24 g·L^-1^ of PHA, which would be in the lower range of previously reported product concentrations.

### Optimization of biofilm formation by *Kyrpidia spormannii* growing on a cathode

Three factors were considered in this study for the optimization of biofilm formation on the cathode: the cathode potential, the oxygen concentration of the sparging of the media, and the pH of the buffered media. The initial conditions were a potential of −525mV, O_2_ concentration of 5% and a pH of 5.5. The current consumption was recorded over 2.8 days, with a plateau after 1 to 2 days, allowing to calculate a stabilized current value for further consideration. The Fig S2-C), with R^2^ at 0.22, 0.47, 0.79 and 0.70 on samples after inoculation, 24 h, 48 h, and 72 h respectively. The average ratio PHA/qPCR were 16.0, 10.1, 6.9 and 4.1 μg·cell^-1^, at 0 h, 24 h, 48 h and 72 h respectively, indicating a divergence of energy into cell multiplication rather than to PHA production during the course of the culture.

Concerning the PHA production, Kourmentza et al. (2017) report PHA concentrations between 0.08 to 2.7 g·L^-1^ (based on reactor volume) produced by different strains using organic carbon sources. Comparatively, our production of PHA is relatively low (maximum of 29.3 ± 1.2 mg·L^-1^), and could probably be optimized by nutrient limitation, as previously described. Up to 90% of the dry cell mass can be composed of PHA (Verlinden et al., 2007). In our case, assuming a mass of 10^-12^ g·cell^-1^, we could theoretically reach 0.24 g·L^-1^ of PHA, which would be in the lower range of previously reported product concentrations.

#### Optimization of cathode potential

The potential screening exhibit two different behaviours over two separate range of potentials (Figure 1, dark-green histograms in the first horizontal panel, Supplementary figure 3). At the most positive potentials, from −325 to −525 mV vs. SHE, no clear trend is observed with current density around 0.03 mA·cm^-2^, while at lower potential, we can see an exponential increase of the maximum current (R^2^=0.975), from 0.44 mA·cm^-2^ at −625 mV vs SHE to 3.77 mA·cm^-2^ at −1425 mV. This increase of current while decreasing the potential is expected by the abiotic reduction of the oxygen on the graphite electrode, with standard potential at 60°C and pH 5.5 estimated at 1.10 V vs. SHE (according to coefficients in Bratsch, 1989). It is then difficult to dissociate the abiotic reaction to the biotic activity of the biofilm. However, the lag-time (Figure 1, light-green histograms in the first horizontal panel) to reach this maximum current is a proxy of the biofilm growth. Indeed, the system is at equilibrium when inoculating the reactor, then, the only increase of current expected is due to biofilm formation, observed by microscopy. This lag time increases to around 0,37 days between −725 and −1025 mV vs. SHE, with a peak at 0,65 ± 0,36 days for −625 mV vs. SHE.

**Figure 1:**
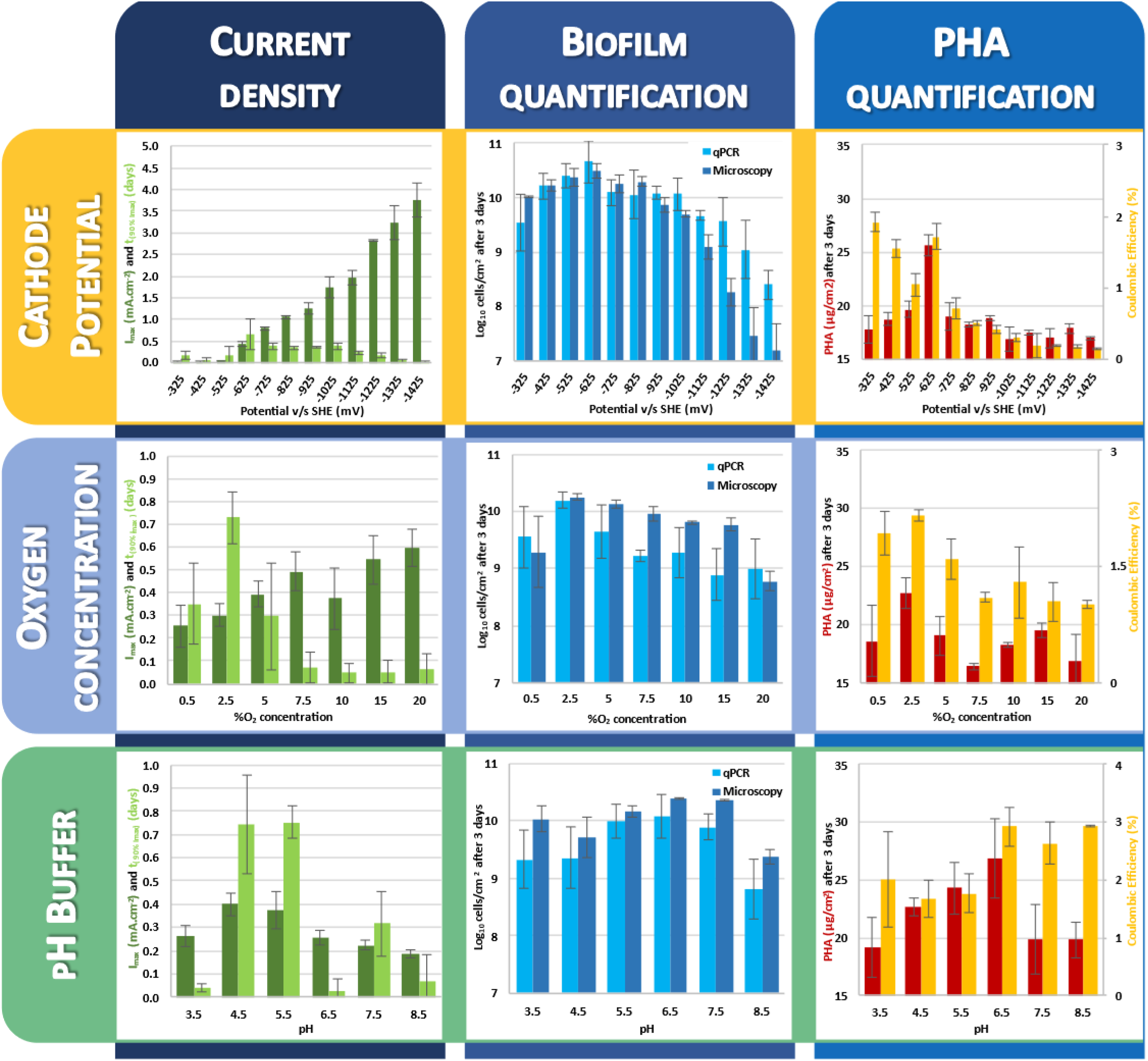
Results of current consumption, lag time, microscopic and qPCR quantifications, PHA production and coulombic efficiency for the optimization of the cathode potential, the oxygen concentration in the MES and the pH of the buffered media. The data presented here are the results of experimental triplicates.

The quantification of the biofilm at the end of the experiment (Figure 1 and 2) indicates a preference for more positive potential, with an increase from 9.5 Log_10_cells·cm^-2^ of electrode at −325 mV vs. SHE to the maximum of 10.5 at −625 mV vs. SHE, followed by a decrease down to 8.4 Log_10_cells·cm^-2^ at −1425 mV vs. SHE on microscopic cell counting. A slight deviation of quantification is observed with the qPCR method, with higher values, probably indicating the death of a part of the biofilm at low potential, not observed in microscopy, but which DNA remains attached to the electrode and quantified by qPCR. This would be corroborated by the known production of toxic H_2_O_2_ or other radicals from two-electron oxygen reduction at low potentials (Pang et al., 2020). Indeed, the H_2_O_2_ production on graphite material was previously reported between −900 to −400 mV vs. SHE in pure oxygen atmosphere, with faradaic efficiency decreasing from 80% to 25% when the potential is more negative (Da Pozzo et al., 2005). However, considering the increase of current at low potential, the total amount of H_2_O_2_ is expected to be higher than at more positive potential, leading to increased death of cells in the biofilm.

**Figure 2:**
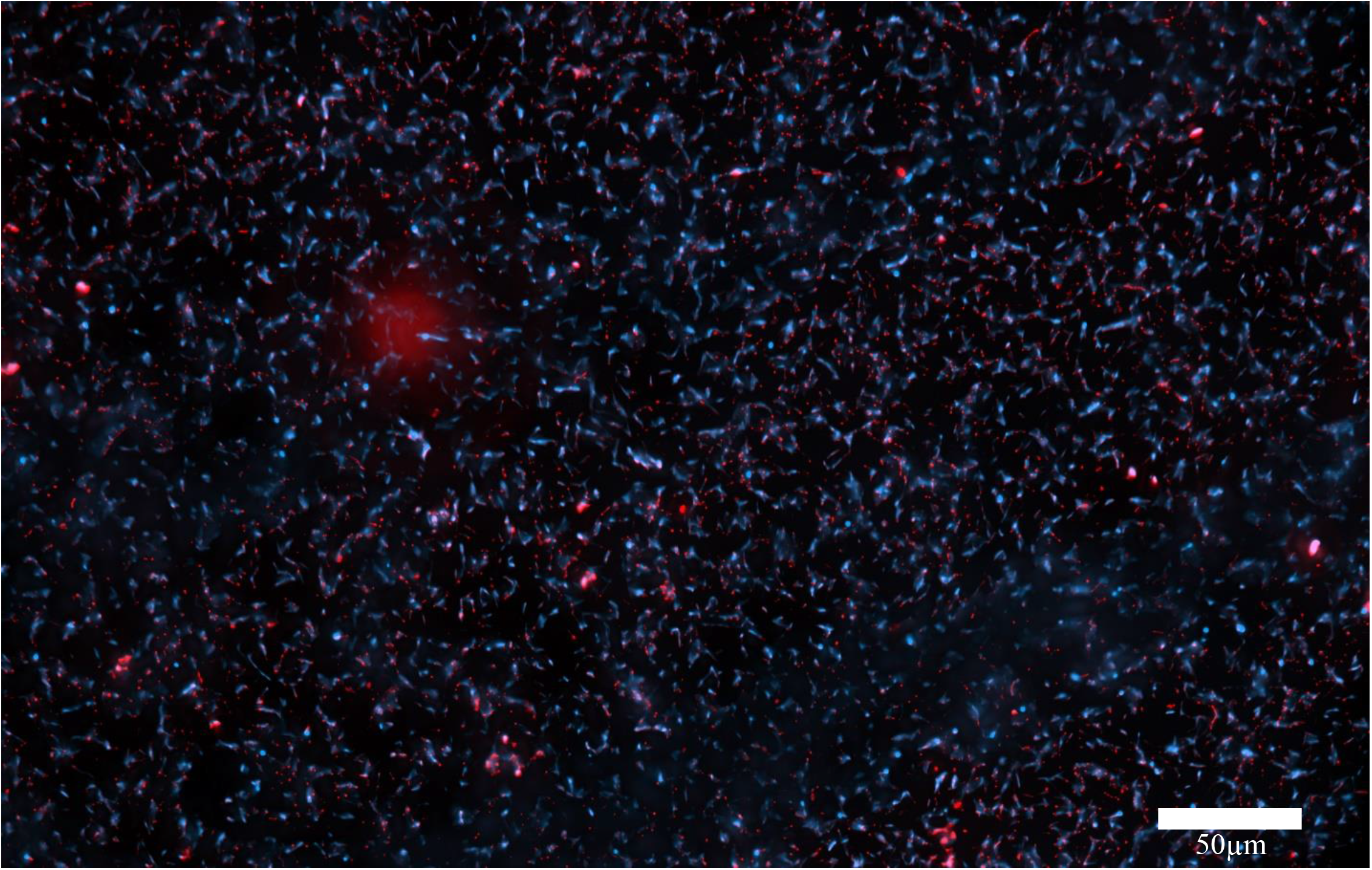
Example of microscopic observation of *Kyrpidia spormannii* EA-1 biofilm on the cathode surface. The blue signal represents the staining of DNA with DAPI, the red signal represents the staining of PHA with NileRed.

A similar effect in observed on the PHA production, with increase from 17.8 to 25.6 μg·cm^-2^ between −325 to −625 mV, followed by a decrease down to 17.0 μg·cm^-2^ at −1425 mV. The coulombic efficiency was calculated at 1.9% at −325 mV decreasing to 0.2% at −1425 mV, with a peak up to 1.7 % at −625 mV. This effect can be explained by the abiotic reaction of oxygen at lower potential, diverging electrons from the cathode to the formation of H_2_O or H_2_O_2_ molecules.

The overall PHA production is relatively low compared to the cells attached to the electrode, with a ratio at only 9.0·10^-9^ μg·cell^-1^ (versus 4.1 μg·cell^-1^ in liquid media). This could be explained by an insufficient detachment step of the PHA by sonication of the biofilm, but microscopy of the electrode after sonication didn’t show any signal with Nile Red staining. It could also be explained by the higher availability of electron donor and acceptor in the MES, with constant electron flow from the cathode and oxygen flow from the gas bubbling. Indeed, in presence of sufficient electron donor and acceptor, the cells proliferate and doesn’t accumulate much PHA. According to the cell concentration and assuming a yield of 90% of the dry mass as PHA, we could theoretically reach 3.4 mg of PHA per cm2 of cathode (versus 25.6 μg·cm^-2^ in our conditions). The potential of −625 mV vs. SHE was then selected for further experiments, as it presented the best biofilm growth, the highest PHA production and one of the highest coulombic efficiency.

#### Optimization of oxygen concentration

Looking at the oxygen effect on biofilm growth and PHA production, presented Figure 1, we can see an overall trend with the increase of current consumption from 0.25 mA·cm^-2^ to 0.59 mA·cm^-2^ while increasing the oxygen concentration from 0.5% up to 20%. However, the lag time (Figure 1, light-green histograms in the second horizontal panel) to achieve this maximum current consumption present a bell curve with a maximum of 0.73 days at 2.5%. At concentration higher than 5%, this delay is reduced to around 0.06 days. This really short delay (1.4 h) is most likely not the result of a microbial growth as it is shorter that the optimal generation time of 3.6 h reported for *Kyrpidia* strains (Hogendoorn et al., 2020) on H_2_:CO_2_. Thus, we can assume that most part of this current consumption is due to abiotic oxygen reduction, especially when increasing O_2_ concentration. The quantification of the biofilm exhibits a similar trend, both in microscopic or qPCR quantification, with maximum biofilm density observed at 2.5% with around 10.2 Log_10_cells·cm^-2^, decreasing down to 8.7-9.0 Log_10_cells·cm^-2^ at 20%. Looking at the PHA production, an optimum of 22.7 μg·cm^-2^ is also observed at 2.5% O_2_, with higher CE up to 2.17%. These values decrease to 16.8 μg·cm^-2^ and 1.02% at 20% O_2_. No increase of PHA production was observed at lower concentration, as expected by the limitation of electron acceptor previously reported in other PHA producers (Kourmentza et al., 2017). Then we can conclude that the optimal O_2_ concentration for *K. spormanni* EA-1 is 2.5% amongst the tested conditions in this work, in agreement with the microaerophilic preference previously reported (Reiner et al., 2018a). The optimal O_2_ concentration for *Kyrpidia* strains in liquid culture is still unknown, but similar O_2_ optimum of 2.5% was observed in other Knallgas bacteria, such as *Mycobacterium genavense* (Realini et al., 1998).

#### Optimization of pH of buffered media

Once the optimal potential and O_2_ concentration were identified, the effect of the pH of the media was studied. As *Kyrpidia* was described as acidophilic, the pH was tested between 3.5 and 8.5. Results associated are presented Figure 1. Looking at the current consumption, a bell curve shape is observed with a maximum at 0.40 mA·cm^-2^ at pH 4.5, decreasing down to 0.19 mA·cm^-2^ at pH 8.5. The delay before stabilization of the current was however more chaotic, with high values around 0.74 days at pH 4.5 and 5.5, intermediate values of 0.32 days at pH 7.5, and values below 0.07 days at pH 3.5, 6.5 and 8.5. The biofilm quantification shows variation of only 1 Log10 between the different pH, with optimum of 10.1-10.4 Log_10_cells·cm^-2^ at pH 6.5, decreasing at 8.8-9.4 Log_10_cells·cm^-2^ at pH 8.5. Finally, the PHA quantification exhibit also a maximum at pH 6.5 with 26.8 μg·cm^-2^ produced with a CE of 2.93%. The PHA production decrease slowly to 19.2 μg·cm^-2^ when decreasing the pH to 3.5 and quickly to 19.8 μg·cm^-2^ when increasing the pH to 7.5-8.5. Thus, an optimum biofilm growth and PHA production is observed at pH 6.5.

The production rate obtained after optimizing the growth conditions reached 96 mg·day^-1^·m^-2^. This value remains relatively low compared to industrial production of PHA from feedstock. However, as previously mentioned, any substrate limitation step was applied here, as the main goal of this work was to produce a dense biofilm prior to this PHA accumulation phase. Further work on substrate limitation of the formed biofilm will allow to more accurately evaluate the industrial potentiality of this new technology.

## CONCLUSION

This study aimed at identifying the optimal conditions for the growth of *Kyrpidia spormannii* EA-1 either in liquid preculture or on the cathode of a Microbial Electrosynthesis System. The results allowed to reduce the culture time from 7 days to 48h by optimizing the substrate, the incubation condition and the media preparation. These results are particularly relevant for the synthesis of PHA in liquid media through lithoauto- or hetero-trophy. The growth of the biofilm was optimized and shows maximum growth of 10.4 Log_10_cells·cm^-2^ and PHA production of 26.8 μg·cm^-2^ or 96 mg·day^-1^om’^2^ at −625 mV vs. SHE, 2.5% O_2_ atmosphere, and a pH of 6.5. These conditions are a starting point to study the effect of nutrient limitation on the formed biofilm for the PHA accumulation in future works. Also, we expect that further optimization of the cathode material and surface modification could increase the initial biofilm growth and PHA production. Only after these optimizations, a meaningful evaluation of the competitiveness of this process for the industrial production of PHA, compared to the heterotrophic or hydrogenotrophic pathways of other PHA producers, will be possible.

## ACKNOWLEDGEMENTS

We are grateful for the financial support from the German Ministry of Education and Research (BMBF) under the program 033RC006B.

## SUPPLEMENTARY INFORMATION

**Supplementary Figure 1:**
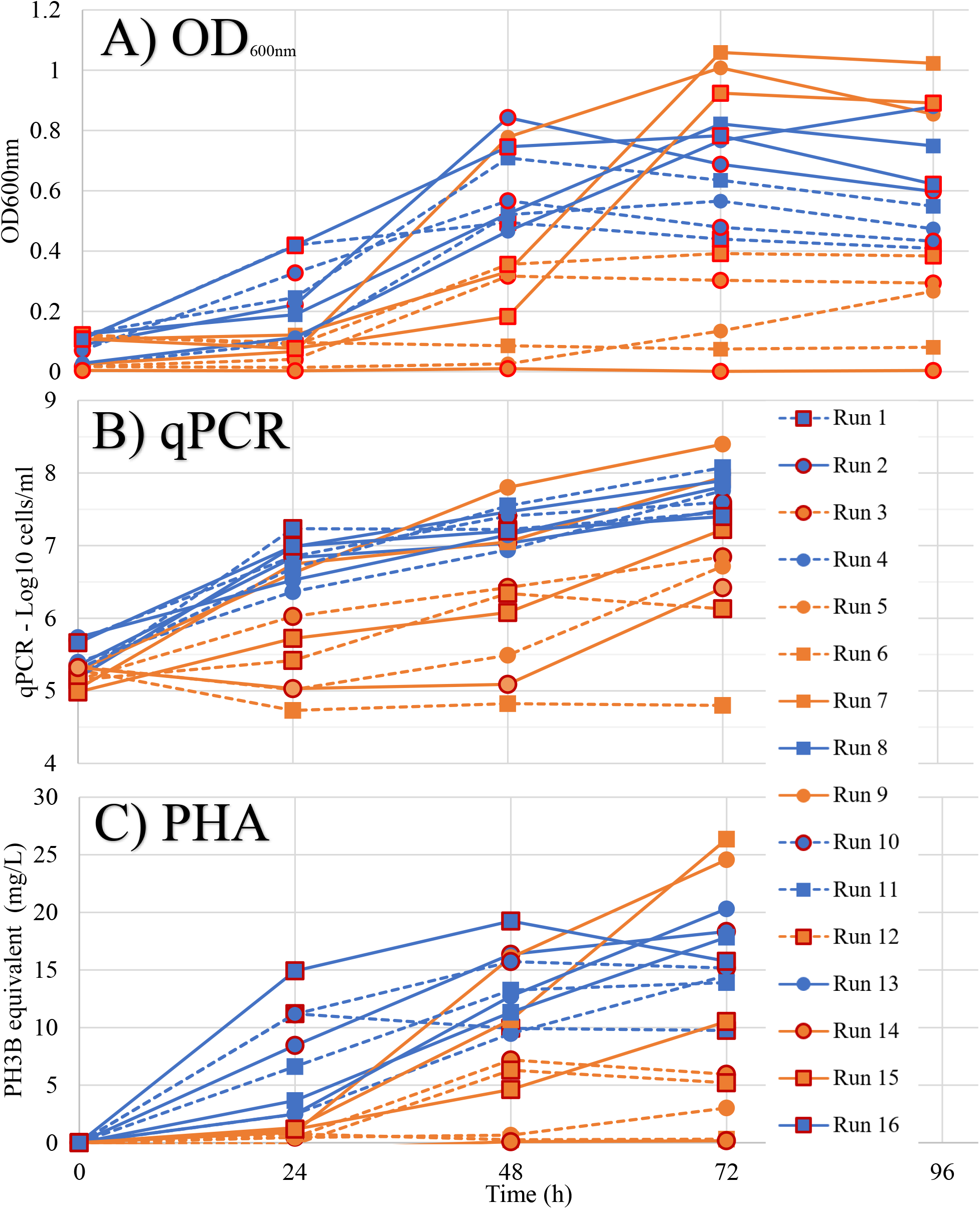
A) average OD_600nm_, B) average qPCR quantification and C) average PHA quantification of triplicates obtained in the 16 runs of the Design of Experiment. Red contour = Shaking; No contour = no Shaking; Blue lines = H_2_:CO_2_; Orange lines = Acetate; Squares = N_2_ media preparation, Circles = O_2_ media preparation; Plain lines = ratio 25/85; Dot lines = ratio 50/60.

**Supplementary Table 1:** Coefficients table of each factor and interaction included in the ANOVA model, with associated p-value. Values in red are significant model terms.

|  | Intercept | A | B | C | D | AB | AC | AD | BC | BD | CD | ABC | ABD | ACD |
| --- | --- | --- | --- | --- | --- | --- | --- | --- | --- | --- | --- | --- | --- | --- |
| OD24h | 0.0908125 | -<br>**0.0425625** | -<br>**0.0919375** | 0.0013125 |  | **0.0526875** |  |  | -<br>**0.0179375** |  |  |  |  |  |
| p-values |  | **< 0.0001** | **< 0.0001** | 0.8284 |  | **< 0.0001** |  |  | **0.0124** |  |  |  |  |  |
| OD48h | 0.348062 | -<br>0.0129375 | -**0.176687** |  | -<br>0.0379375 | 0.0310625 |  | -<br>0.0309375 |  | -<br>0.0051875 |  |  | -<br>**0.125437** |  |
| p-values |  | 0.6097 | **< 0.0001** |  | 0.1579 | 0.2379 |  | 0.2396 |  | 0.8366 |  |  | **0.0009** |  |
| OD72h | 0.487 | **0.076125** | **-0.07625** |  | **-0.183375** | 0.016375 |  | **-0.09625** |  | **-0.069125** |  |  | **-0.1155** |  |
| p-values |  | **0.0112** | **0.0111** |  | **< 0.0001** | 0.5007 |  | **0.0032** |  | **0.0177** |  |  | **0.0011** |  |
| qPCR24h | 6.33568 | -0.12605 | **-0.473542** | -<br>0.0211864 | **-0.294625** |  |  | **-0.214192** | **-0.186941** | **-0.270464** |  |  |  |  |
| p-values |  | 0.1022 | **0.0001** | 0.7643 | **0.0026** |  |  | **0.0139** | **0.0256** | **0.0042** |  |  |  |  |
| qPCR48h | 6.86071 | -0.077208 | **-0.384647** | -0.14466 | **-0.373705** |  |  | **-0.208833** | **-0.258029** | **-0.407826** |  |  |  |  |
| p-values |  | 0.4015 | **0.0022** | 0.1355 | **0.0027** |  |  | **0.0434** | **0.0181** | **0.0016** |  |  |  |  |
| qPCR72h | 7.43325 |  | **-0.252373** | **-0.193156** | **-0.38614** |  |  |  | **-0.214765** | **-0.422631** |  |  |  |  |
| p-values |  |  | **0.0035** | **0.0156** | **0.0002** |  |  |  | **0.0089** | **< 0.0001** |  |  |  |  |
| PHA24h | 4.08385 | **-1.75164** | **-3.53669** | **0.853453** | 0.0444391 | **2.05968** | -<br>0.102823 | 0.198316 | -0.621802 |  | -<br>0.370175 |  |  | **0.728264** |
| p-values |  | **0.0010** | **< 0.0001** | **0.0210** | 0.8696 | **0.0005** | 0.7058 | 0.4755 | 0.0603 |  | 0.2096 |  |  | **0.0366** |
| PHA48h | 9.61721 | -0.314248 | **-3.88611** | -0.168951 | **-1.77043** | **1.49753** | -<br>0.259901 | **-1.62439** | -0.102226 | -0.360546 | -<br>0.243937 | -<br>**0.920245** | **-2.7015** | **1.51758** |
| p-values |  | 0.2639 | **0.0028** | 0.4953 | **0.0131** | **0.0181** | 0.3313 | **0.0155** | 0.6665 | 0.2197 | 0.3549 | **0.0459** | **0.0057** | **0.0176** |
| PHA72h | 12.6036 | **2.49875** | **-3.09133** |  | **-4.12745** | 1.55219 |  | **-3.03736** |  | -1.76443 |  |  | **-2.96805** |  |
| p-values |  | **0.0135** | **0.0045** |  | **0.0008** | 0.0858 |  | **0.0050** |  | 0.0566 |  |  | **0.0057** |  |

**Supplementary Figure 2:**
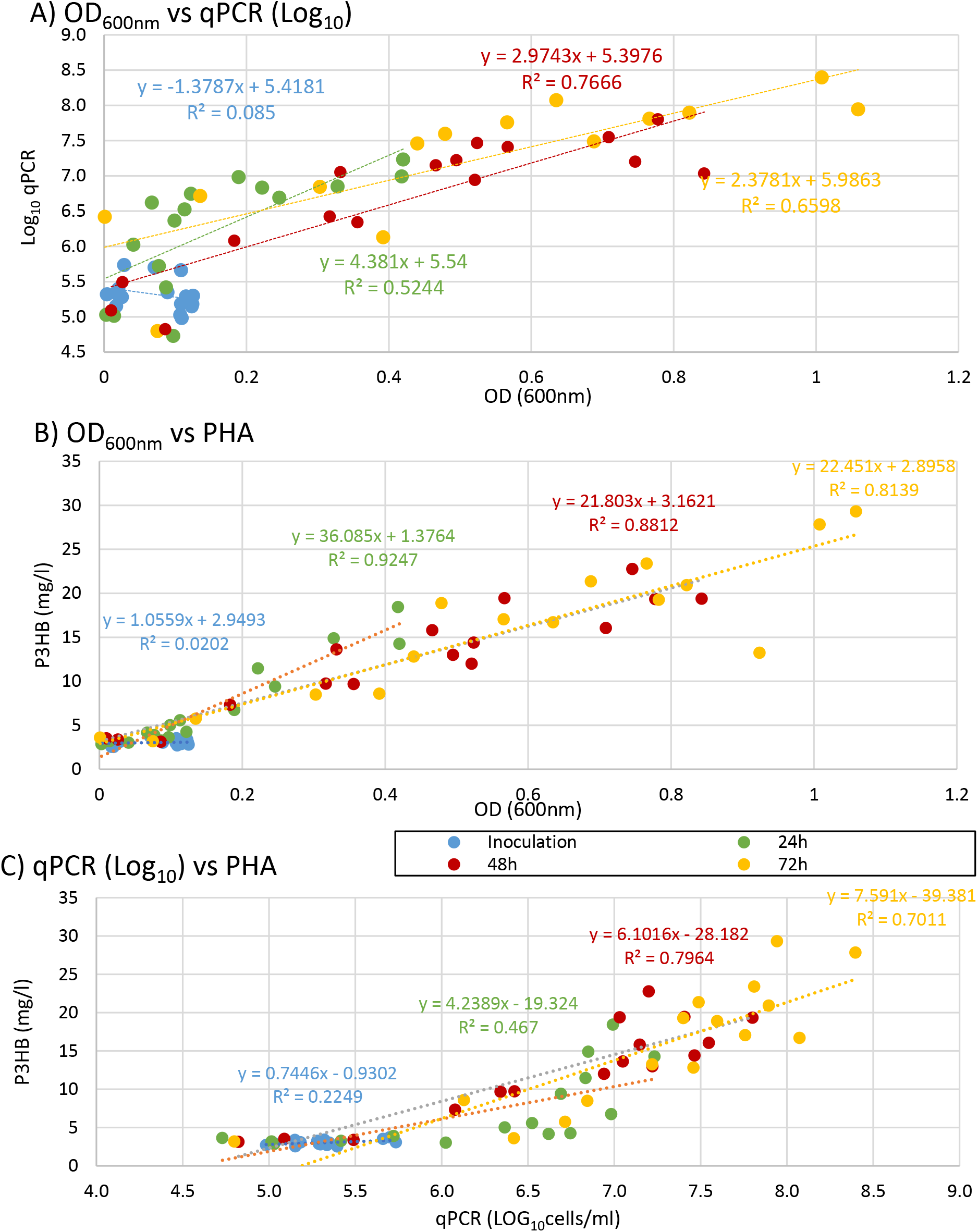
Correlation between A) OD_600nm_ measurements and qPCR quantification, B) OD_600nm_ measurements and PHA production and C) qPCR measurement and PHA production, in *Kyrpidia spormannii*.

**Supplementary figure 3:**
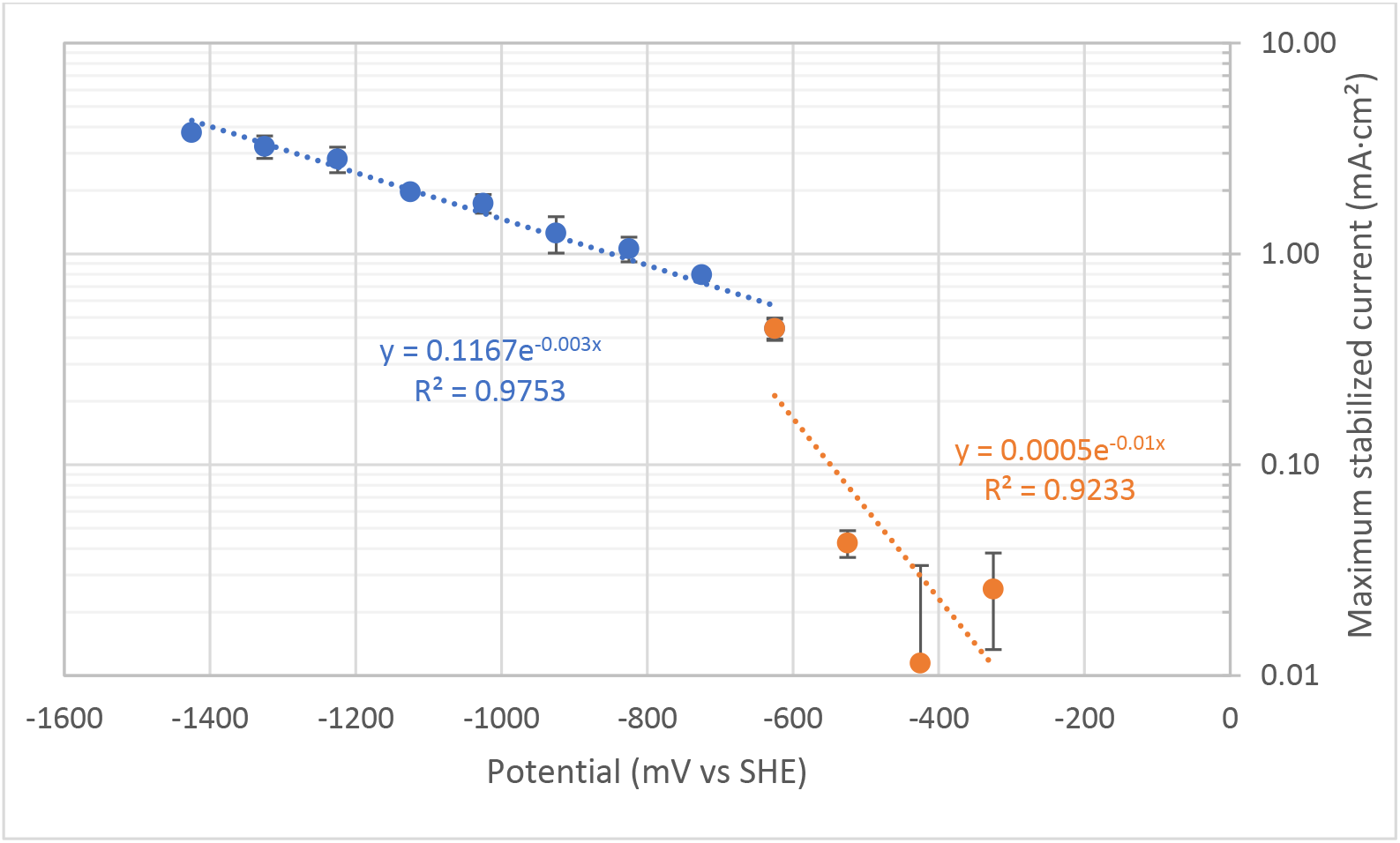
Exponential regression of the maximum current obtained vs. potential of the cathode. Two separate trends can be observed, here represented by the blue and orange colours.

